# Intraspecific scaling of home range size and its bioenergetic dependence

**DOI:** 10.1101/2023.06.08.544203

**Authors:** Evan E. Byrnes, Jenna L. Hounslow, Vital Heim, Clemency E. White, Matthew J. Smukall, Stephen J. Beatty, Adrian C. Gleiss

## Abstract

Home range size and metabolic rate of animals are expected to scale with body mass at similar rates; with home ranges expanding to meet increased metabolic requirements. This expectation has widely been tested using lab-derived estimates of basal metabolic rate as proxies for field energy requirements, however, it is unclear if existing theory aligns with patterns of home range scaling observed in the field. Here, we conduct the first direct field test of the relationship between home range and metabolic rate allometry. Using acoustic telemetry, we simultaneously measured the individual home range size and field metabolic rate of lemon sharks *(Negaprion brevirostris)* spanning one order of magnitude in body mass. Although scaling rates of field metabolic rate were consistent with standard metabolic rate, home range size scaled at shallower rates than metabolic rates. This is evidence for strong top-down controls on home range scaling rates, likely a result of predation pressure placing constraints on home range expansions. Consequently, direct resource competition can lead to decreased home range scaling rates. We highlight inconsistencies with theory on the effects of population density and competition on home range scaling and propose that the influence of diverse types of competition should be examined.

## INTRODUCTION

With the exception of nomads and migrants, animals tend to move within a restricted home range, which is widely observed to increase allometrically with body size (Teitelbaum and Mueller 2019). The size of home ranges has long been thought to scale proportionately with the metabolic rate of animals, as their home range must provide access to sufficient resources for sustenance. To test this idea, McNab (1963) estimated the allometric scaling rates of interspecific mammalian basal metabolic rate (BMR) and home range size and found both scaled at a similar rate. This finding led to the assertation that home range size is directly proportional to metabolic requirements. Subsequent field studies, however, found that home range size scaled at substantially higher rates than predicted by McNab’s hypothesis (e.g., Lindstedt et al. 1986, Minns 1995, Pearce et al. 2013). The most prominent explanation for such discrepancies is that animals with overlapping home ranges share resources, and as animals grow, the proportion of resources that are shared and the costs of defending them increases (Damuth 1981, Jetz et al. 2004). Contrary to the assumptions of this explanation, not all species actively defend territories or resources but rather share resources through scramble competition. Accordingly, field studies have presented conflicting support for this shared resource hypothesis (e.g., Pearce et al. 2013, Ofstad et al. 2016).

One potential, yet often overlooked explanation for home range size scaling at steeper rates than predicted by existing theory is that the field metabolic rates of animals are higher than assumed in current models. Various methodological constraints have limited our ability to estimate metabolic rate over long time scales (see Butler et al. 2004, Wilson et al. 2020), making it previously impossible to directly compare allometries of FMR and home range. As a result, current hypotheses on the relationship between metabolic rate and home range size were built using estimates of BMR of animals (McNab 1963, Haskell et al. 2002, Jetz et al. 2004), and therefore assume that BMR and field metabolic rates (FMR) are synonymous. However, this is unrealistic given that FMR of animals include costs associated with foraging and evading predators, growth, and in mature individuals, reproduction, all of which may respond to different ecological circumstances. Therefore, animals require more resources and in consequence, more space than predicted based on BMR alone. Empirical results across various taxa have shown active (i.e., routine) metabolic rates scale allometrically at higher rates than maintenance (i.e., basal or standard) metabolic rates (Weibel et al. 2004, Killen et al. 2010). Therefore, observations where home range size scaled at higher rates than predicted by McNab (1963) may simply be a product of animals attempting to acquire resources required to meet active metabolic rate (i.e., FMR).

Clarifying the relationship between metabolic rate and home range size has been challenging because factors other than metabolism can simultaneously influence home range size. These factor include locomotory mode and speed, thermoregulation strategy, foraging niche and dimensionality, and social organization (Tamburello et al. 2015, Papageorgiou and Farine 2020). The influence of these mechanisms on home range size inherently varies between taxa. For example, animals that forage at higher trophic levels tend to experience lower resource densities, and thus may require larger areas than animals with a similar metabolic rate that forage on a lower trophic level (Tamburello et al. 2015). Nevertheless, McNab’s bioenergetic theory of home range allometry has only been tested with interspecific studies (e.g., Jetz et al. 2004, Haskell et al. 2002, Noonan et al. 2020), making it difficult to discern to the impact of metabolism from these other mechanisms.

Disentangling the various drivers of home range size is particularly difficult in higher vertebrates. In birds and mammals, parental investment and complex social systems impact metabolic demands (Alonso-Alvarez and Velando 2012, Pearce et al. 2013), confounding the metabolic dependency of individual home range size across different life stages. However, in many lower vertebrates these mechanisms can be more easily separated. For example, elasmobranchs are a useful model because they grow by several orders of magnitude over ontogeny and are self-sufficient foragers throughout all life stages, enabling us to investigate home range allometry independent of unsystematic differences in metabolic demands.

In this study, we address two key questions about the spatial ecology of animals by quantifying the extent of home range size and FMR of lemon sharks (*Negaprion brevirostris*) in tandem. Importantly, we produce the first estimates of FMR across an order of magnitude of body-mass using the acceleration method (Gleiss et al. 2011) and comprehensively validated our metabolic proxy across the same scales of body mass (Byrnes et al. 2021). From these novel estimates of FMR, we first test whether the scaling of home range size and FMR conform to predictions made by existing theory by directly estimating these variables for individuals spanning one order of magnitude in body size. Second, we explore potential mechanisms underpinning the observed scaling rates. Through the early life stages, lemon sharks reside in nursery habitats to mitigate high risk of mortality from predation (Morrissey and Gruber 1993). However, the protection provided by the spatial restrictions of the nursery area is accompanied by heavy resource competition, caused by seasonal influxes of neonates, which provides an annual oversupply of competitors (Feldheim et al. 2002, Dibattista et al. 2007). As such, in addition to metabolic rate, our system allows insight on the function of predation pressure and resource competition in determining home range allometric scaling rates.

## METHODS

### Study site and species

The study was conducted at Bimini, Bahamas (25°44’N, 79°16’W), a mangrove-fringed chain of islands located approximately 85 km east of Miami, Florida, USA. The Bimini Islands enclose an approximately 21 km^2^ lagoon that is between 0-1.2 m deep at low tide. The relatively shallow water depth limits the abundance of large marine predators, providing a nursery habitat for juvenile lemon sharks (Morrissey and Gruber 1993, Heupel et al. 2007). However, body size dependent predation risk is known to place large constraints on habitat use within the nursery, causing different life stages to partition between different available habitats (e.g., mangrove channels, shallow sand flats, Dhellemmes et al. 2021b). Consequently, high numbers of younger individuals tend to concentrate into small areas of habitat, leading to inflated levels of resource competition, indicated by high rates of mortality within the first few years of life (Gruber et al. 2001, Dibattista et al. 2007). Moreover, adult females return to the nursery to give birth each spring, causing an annual influx of additional competitors for early life stages (Feldheim et al. 2002). Individuals show high site fidelity to their pupping area through at least the first three years of life (Morrissey and Gruber 1993), after which home range size increases and sharks gradually disperse into deeper habitats around the lagoon, with emigration near sexual maturity (∼2.3 m total length, 2009). This high site fidelity through the early life stages allows for home range size and daily metabolic requirements to be reliably quantified over long periods of time for individuals spanning a continuum of body sizes. Additionally, focusing on juveniles allowed us to preclude the influence of reproduction on spatial behaviours (e.g., migrations) or energy costs (e.g., somatic growth).

### Tagging and data preparation

From April to August 2019, 20 lemon sharks were captured in the lagoon between North and South Bimini using a combination of handline, drumline, and long-line fishing methods. Upon capture, sharks were sexed, measured for length (precaudal (PCL), fork (FL), and total (TL)) and implanted with an acoustic tag (V13AP; Vemco, Innovasea, NS, CAN). Tags were surgically implanted in the peritoneal cavity of each shark through a 4 mm incision that was sealed with two simple interrupted sutures using poliglecaprone 25 sutures (Q310 MonoWeb, Patterson Veterinary, Devens, MA, USA). As part of additional studies, one fin clip, a muscle biopsy, and blood sample were also taken from each shark prior to release. This entire sampling and tagging procedure lasted between 10-15 minutes. After release at the site of capture, tags alternately transmitted body acceleration (m/s^2^) and depth (m) data at a nominal delay of 90-180 seconds. Body acceleration data were recorded at 5 Hz for a period of 20 seconds, which was processed onboard tags as mean vectorial dynamic body acceleration (VeDBA) over the entire recording duration. Tag transmissions were recorded by a network of 60 underwater receivers placed around Bimini (Fig. 1). Temperature loggers (HOBO U22-001 and HOBO MX2201; Onset Computer Corp., MA, USA) were attached to 40 receivers, representing all available habitats and recorded ambient water temperature every 10 min. In December 2019 and January 2020, receivers and temperature loggers were retrieved to download acoustic detections and temperature data. All animal use was conducted in accordance with permits from the Bahamas Department of Marine Resources (MA&MR/FIS/178) and Murdoch University Animal Ethics committee (RW3119/19). Prior to analysis, the first week of data for all tags were removed to allow sharks to recover from the tagging procedure and all double and false detections were removed from the dataset as per Kessel *et al*. (Kessel et al. 2014). Additionally, to ensure that home range and metabolic rate estimations were made over the same timeframe for all sharks, only data during which all tags were active (July 28^th^ through December 12^th^, 2019) were used for analyses.

**Fig 1.**
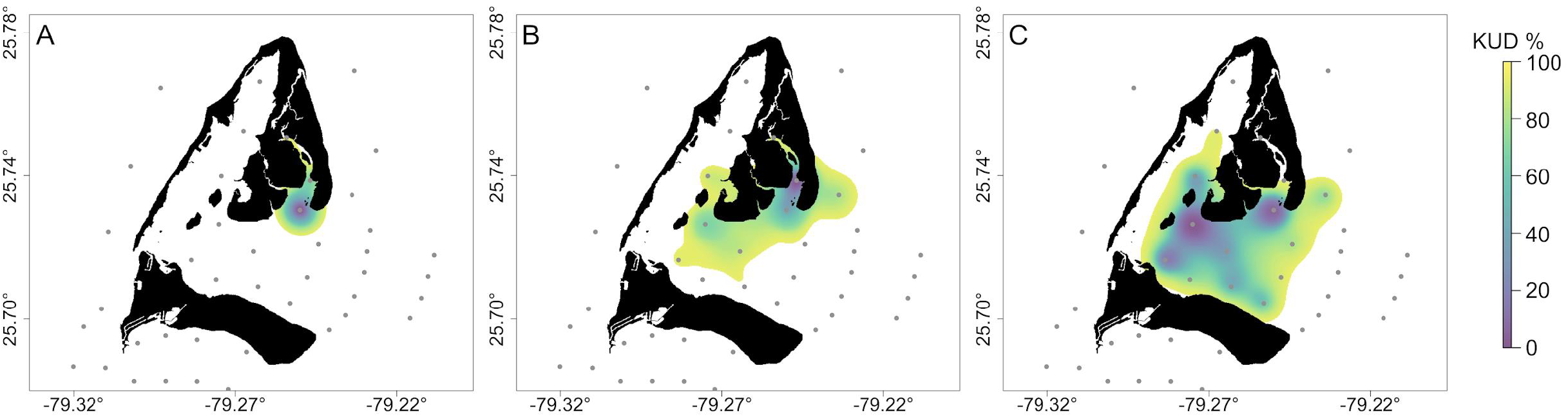
Home ranges of three lemon sharks (*Negaprion brevirostris)*, in Bimini, BHS, demonstrating how home range size increased with body size. For clarity, only three sharks are displayed, representing the smallest (left; 2.32 kg and 1.82 km^2^), a mid-sized (middle; 9.74 kg & 9.38 km^2^), and the largest (right; 17.76 kg & 16.71 km^2^) individuals. Receiver locations are indicated by grey dots; not all 60 receivers included in figure. Home ranges for all tagged individuals provided in Table 1.

### Home range analysis

Home range size (95% utilization distribution) was calculated for each individual using functions provided in the ‘Animal Tracking Toolbox’ extension of the ‘VTrack’ package (Udyawer et al. 2018) in the R statistical environment, (version 3.6.3, R Core Team 2019). For preparation of home range analysis, three-hourly mean geographic positions were estimated for each shark using the ‘COA’ function. The use of mean geographic position estimates in home range analysis, rather than raw locations provides a more accurate representation of animal movement by accounting for temporally variable tag transmissions and spatial biases from fixed receiver locations (Simpfendorfer, Heupel & Hueter 2002).

To estimate the individual home range sizes, Brownian bridge movement models (BBMMs) were applied to centre-of-activity estimates from each shark using the ‘HRsummary’ function. BBMMs require input of two initial parameters: 1) the Brownian motion variance (σ^2^_m_), representing how diffusive or irregular the movement of an animal is and 2) the error associated with location estimates. The σ^2^_m_was estimated within the ‘HRsummary’ function, which uses the likelihood maximization method established by Kranstauber et al. (2012). While the ‘HRsummary’ function normally requires at least six locations to estimate σ^2^_m_, the limited spatial movement demonstrated by some sharks did not provide sufficient data to meet these requirements. However, the approach of Kranstauber et al. (2012) utilizes a “leave-one-out” method, which only requires three locations to estimate σ^2^_m_. Therefore, we adjusted the requirements within the ‘HRsummary’ function to a minimum of three locations to ensure home ranges could be estimated for all sharks within the study. Location error was set to 255 m, which is equal to the detection range estimated for receivers at the study site (Guttridge et al. 2017). BBMMs were used to estimate 95% utilization distributions for each individual for the entire study period. Land was excluded from utilization distributions by overlaying them on a habitat raster. Areas overlapping land were manually clipped and the remaining utilization distribution was estimated to the nearest 1 m^2^.

### FMR estimation

To estimate daily energy requirements of each shark, mean daily FMR was back calculated using the bioenergetic equation:

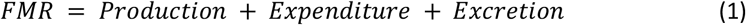

Production was calculated based on estimated growth rates of each shark. Growth rates were estimated using the von Bertalanffy growth curve established for lemon sharks (Brown and Gruber 1988):

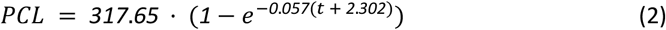

where PCL is precaudal length at the time of capture and *t* is the estimated age of the fish at time of capture. First, the age of the shark upon capture was estimated by inserting the precaudal length at capture into equation 2. Then one year was added to the age at capture and a presumptive precaudal length was estimated using equation 2. Mass of the shark at time of capture and one year later were estimated based on exponential length to weight relationships established from sharks captured at Bimini as part of other experiments (*R*^*2*^ = 0.85, *N* = 399; unpublished data). The difference between these masses was converted to a daily energy equivalent by multiplying by the energy content of a lemon shark tissue, 5.4 kJ g^-1^ (Cortes and Gruber 1994) and dividing by 365 days yr^-1^.

Oxygen uptake rates (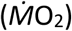) were estimated as proxies of FMR, based on calibrated equations relating 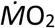 and VeDBA. Prior to 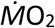 estimation, VeDBA values were corrected to account for acceleration sensor noise; sensors at complete rest record small acceleration values that would inflate VeDBA estimates. To correct values, raw VeDBA observations were labelled as either active (i.e., swimming) or inactive (i.e., resting) using histogram segregation, with higher values indicating active and lower values indicating inactive behaviour (Fig. S1, Collins et al. 2015). Mean sensor noise was then estimated as the mean VeDBA recorded during inactive observations, which was then subtracted from all active VeDBA observations. Inactive VeDBA observations were all set to equal zero, as these should represent periods when animals were resting motionless on the bottom. 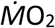 were predicted for each available detection based on VeDBA and individual mass data using the oxygen uptake rate predictive equation established for this population of lemon sharks by Byrnes et al. (2021):

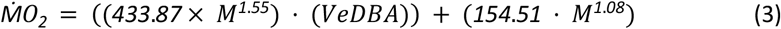

To account for the temperature effects in 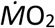 estimations, the intercept was adjusted by correcting 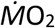 estimates using a Q_10_ relationship (Gillooly et al. 2001):

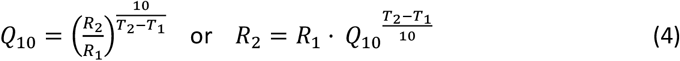

where *Q*_*10*_ is the temperature correction factor of 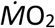, *T*_*1*_ is the temperature at which equation 3 was calibrated (29.50 °C), *T*_*2*_ is the observed water temperature at the time of a respective detection, *R*_*1*_ is the 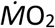 estimated at *T*_*1*_, *R*_*2*_ is the 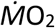 at *T*_*2*._ A *Q*_*10*_ of 2.96 was used for all inactive detections, and a *Q*_*10*_ of 1.69 was used to for all active detections (Lear et al. 2017). Water temperatures used in 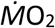 predictive equations were estimated using random forest (RF) regression models (see online supplement). Hourly water temperature was estimated at each receiver location based on a suite of environmental variables (Table S1), which were then time matched to acoustic detections. Overall, RF models for each receiver had a mean squared error ranging from 0.17 -1.41 °C (mean: 0.67 °C; Table S2). Mean daily FMR was calculated for each individual. In addition, since inactive detections were not recorded for all individuals, standard metabolic rate (SMR; analogue of BMR for heterotherms) was estimated by calculating the daily FMR using equation 3, with VeDBA set to zero.

Excretion was assumed to equal 27% of FMR, based on an 80% absorption efficiency (Wetherbee and Gruber 1993) and 7% loss of assimilated energy in gill and urine effluent (Brett and Groves 1979).

### Statistical analysis

The allometry of home range size and daily FMR were explored with least squares regression in the base R ‘stats’ package (R Core Team 2019). Before fitting regressions, body mass, home range size (i.e., overall trimmed 95% kernels) and mean daily FMR estimates were natural log transformed. *ln*(home range size) and *ln*(mean daily FMR) were plotted separately against *ln*(body mass), and confidence intervals of model parameters were determined using the ‘confint’ function, which uses the profile likelihood method. Lastly, final model formulae were exponentiated to establish the power functions describing the allometric scaling of home range size and daily FMR.

## RESULTS

### Deployments

Two individuals left the array of acoustic receivers within 12 days of being tagged, and as such, did not provide sufficient data to estimate home range size. Home range size and daily FMR were estimated for the remaining 18 individuals, ranging in estimated body mass from 2.32 to 17.76 kg (Table 1). One of the 18 sharks was also excluded from further analysis because it demonstrated an uncharacteristically small home range, indicating that it died or shed its tag (Table 1). For the remaining 17 individuals, home range size ranged from 2.04 to 21.35 km^2^; overall daily FMR ranged from 712.83 to 7,358.18 kJ day^-1^, and SMR from

**Table 1.** Summary of biometrics, home range size, FMR, and SMR of all sharks.

| Date Tagged | Sex | Total length (cm) | Mass (kg) | No. of detections | % detections active | No. days detected | Mean temp (°C) | Home range size (km<sup>2</sup>) | Mean FMR (kJ day<sup>-1</sup>) | Mean SMR (kJ day<sup>-1</sup>) |
| --- | --- | --- | --- | --- | --- | --- | --- | --- | --- | --- |
| 6/04/2019 | F | 131.00 | 10.94 | 4,951 | 84.62 | 233 | 27.80 | 1.52<sup>†</sup> | - | - |
| 6/04/2019 | M | 145.00 | 14.83 | 1,515 | 93.40 | 176 | 29.20 | 17.76 | 5,694.13 | 2,678.84 |
| 12/04/2019 | F | 99.00 | 4.73 | 2,807 | 99.90 | 199 | 27.10 | 3.63 | 1,560.167 | 731.00 |
| 12/04/2019 | F | 126.00 | 9.74 | 1,971 | 96.95 | 177 | 28.90 | 9.38 | 3,620.35 | 1,706.64 |
| 20/04/2019 | F | 121.00 | 8.63 | 1,548 | 100.00 | 148 | 28.90 | 16.11 | 3,255.34 | 1,458.90 |
| 21/04/2019 | F | 113.00 | 7.03 | 2,948 | 100.00 | 184 | 28.20 | 2.04 | 2,433.18 | 1,160.48 |
| 5/05/2019 | F | 163.00 | 21.06 | 48 | - | 12 | - | _<sup>‡</sup> | _<sup>‡</sup> | _<sup>‡</sup> |
| 6/05/2019 | M | 102.20 | 5.20 | 380 | 98.44 | 84 | 27.30 | 16.00 | 1,841.95 | 823.95 |
| 6/05/2019 | F | 108.00 | 6.14 | 2,151 | 97.44 | 163 | 28.40 | 5.03 | 2,306.26 | 1,071.68 |
| 6/05/2019 | F | 131.50 | 11.07 | 2,214 | 99.21 | 178 | 28.30 | 21.35 | 4,569.71 | 1,885.80 |
| 8/05/2019 | F | 134.40 | 11.82 | 2,675 | 99.79 | 151 | 29.60 | 7.21 | 4,689.77 | 2,203.44 |
| 16/05/2019 | M | 97.00 | 4.45 | 300 | 94.39 | 50 | 28.80 | 3.34 | 1,531.55 | 715.21 |
| 18/05/2019 | M | 144.50 | 14.68 | 2,361 | 99.89 | 173 | 28.70 | 10.63 | 6,080.72 | 2,673.90 |
| 19/05/2019 | M | 78.00 | 2.32 | 822 | 100.00 | 105 | 29.00 | 1.82 | 712.83 | 374.68 |
| 19/05/2019 | M | 127.00 | 9.97 | 1,311 | 100.00 | 147 | 28.90 | 19.17 | 3,925.03 | 1,786.53 |
| 21/05/2019 | M | 97.50 | 4.52 | 817 | 100.00 | 92 | 26.10 | 9.96 | 1,486.63 | 671.60 |
| 3/06/2019 | M | 179.00 | 27.87 | 13 | - | 1 | - | _<sup>‡</sup> | _<sup>‡</sup> | _<sup>‡</sup> |
| 4/07/2019 | F | 154.00 | 17.76 | 1,374 | 96.71 | 112 | 28.70 | 16.71 | 7,358.18 | 3,382.10 |
| 7/07/2019 | F | 133.00 | 11.45 | 1,744 | 99.87 | 124 | 28.40 | 14.42 | 4,310.59 | 1,990.40 |
| 20/07/2019 | F | 115.00 | 7.41 | 1,184 | 100.00 | 61 | 30.10 | 6.97 | 2,881.82 | 1,424.70 |

<sup>†</sup>Shark was removed from analysis due to an uncharacteristically small home range indicating that the shark died, or the tag was shed prematurely.
<sup>‡</sup>Insufficient data recorded for estimation of home range or metabolic rates.

374.68 to 3,382.10 kJ day^-1^ (Table 1). Sharks were active for the majority of detections, ranging from 93.40% to 100.00%. Overall, temperatures experienced by sharks ranged from 20.20 °C to 41.40 °C. Mean temperatures experienced varied among individuals, ranging from 26.10 °C to 30.10 °C (Table 1). However, there was no evidence that body mass had an influence on mean temperature experienced (*t* = 1.49, *p* = 0.16) (Fig. S2)

### Home range and FMR allometry

Body mass demonstrated a significant positive influence on home range size (*r*^2^ = 0.47; *F*_*1,15*_ = 13.41, *p* < 0.01), daily FMR (*r*^2^ = 0.98; *F*_1,15_ = 1032, *p* < 0.01), and daily mean estimated SMR (*r*^2^ = 0.48; *F*_*1,15*_ = 15.51, *p* < 0.01). Home range size scaled positively with body mass with an allometric exponent of 1.01 (95% CI: 0.42 -1.60) (Fig. 1,2). Daily FMR also scaled positively with body mass with an allometric exponent of 1.15 (95% CI: 1.11 -1.18) (Fig. 2). Lastly, mean daily SMR scaled with a slightly lower allometric exponent than FMR at 1.10 (95% CI: 1.05 -1.15) (Fig. 2).

**Fig 2.**
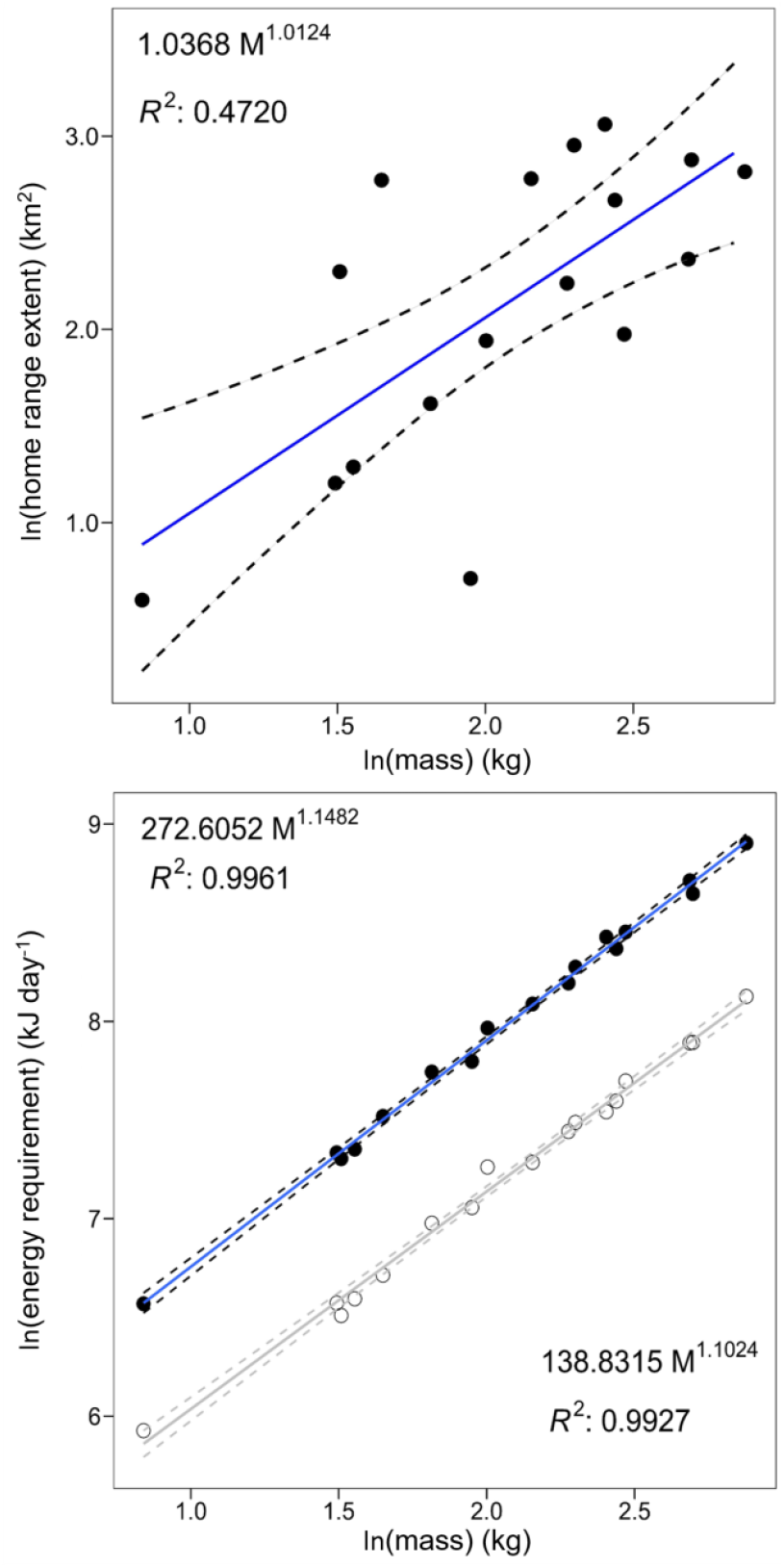
Natural-log linear function between home range size and mean daily FMR with corresponding allometric scaling relationship and *R*^*2*^. For comparison, mean daily SMR (open circles, gray line) is provided plotted along with daily FMR. Allometric scaling relationships were established by exponentiation of the natural logarithm function.

## DISCUSSION

We provide the first direct field test of whether allometric changes in home range size scale proportionally to changes in field energetic requirements, which was made possible by the development of a proxy for metabolic rate explicitly incorporating mass (Byrnes et al. 2021). Meeting metabolic demand is one of the most fundamental drivers of animal ecology, placing large constraints on behaviours and life history of individuals (Brown et al. 2004). As such, it is expected that the home range size of animals directly represents their energy requirements; at a minimum, home range size should scale equally to energy requirements (McNab 1963). However, our observations of a shallower intraspecific scaling rate of home range size (M^1.01^) than FMR (M^1.15^) paradoxically did not match such expectations, indicating that mechanisms influencing home range allometry do not apply universally across different levels of taxonomic organization. Moreover, home range size scaled at shallower rates than SMR (M^1.10^), suggesting that energy requirements alone could not be responsible for determining observed home range sizes. This divergence in home range and metabolic allometries broadly agrees with recent models that suggest several other ecological factors can alter the scaling exponents of home range size (Haskell et al. 2002, Jetz et al. 2004, Carbone et al. 2005). However, these models tend to examine mechanisms that cause inflated home range scaling rates, whereas our results indicate that other mechanisms exist that may limit scaling rates. Furthermore, the scaling rates observed here suggest that these other mechanisms can place stronger constraints on home range size than energy requirements.

It is well established that predation pressure places substantial constraints on the distribution and movements of animals (Preisser and Orrock 2012). While larger animals are afforded a size refuge from predation, smaller animals may rely more heavily on habitat refuges to mitigate risk of mortality (Chase 1999). As such, smaller animals often constrain the distance travelled from a refuge and occupy habitats with high refuge densities compared to larger animals (Crowell et al. 2016). Indeed, many elasmobranchs species (including lemon sharks) limit their movements to small areas of coastal habitat during their early life stages, to take advantage of the protection provided by shallow embayments and fringing vegetation (Heupel et al. 2007). Considering that spatial distributions of smaller lemon sharks (∼ <130 cm TL) change in response to predator presence (Wetherbee et al. 2007, Guttridge et al. 2012), it is therefore also likely that predation risk influences their home range. Since all individuals for this study were at body sizes prone to predation, we believe the reduced home range scaling rates observed were likely a result of the increased predation risk faced by smaller animals.

Home range size scaled at a shallower rate than metabolic rate, which was unexpected given that resource competition is expected to have a positive effect on home range size. In high levels of resource competition, animals will expand their home range to mitigate resource limitations, resulting in home range size scaling at greater rates than energy requirements (Haskell et al. 2002, Jetz et al. 2004). High levels of resource competition are a common characteristic of nursery habitats (Bush and Holland 2002, Heupel et al. 2007), where high juvenile mortality rates have commonly been attributed to starvation in multiple species of coastal elasmobranchs. Indeed, within our study system juvenile lemon sharks have high mortality rates, indicating strong resource limitations attributed to competition (Dibattista et al. 2007). Nonetheless, although we expected home range to scale at a steeper rate than metabolic rate, it is possible that increased competition for resources may not always lead to increased home range scaling rates. For instance, in this system, it is likely that the advantages of home range expansions are negated by top-down pressures that pose a more immediate risk to survival than starvation, such as risk of mortality from predation (Gallagher et al. 2017). Moreover, in highly fragmented habitats, such as archipelagos, reduced access to additional resources patches may also limit the advantages of home range expansions.

Here we also consider how the effect of resource competition on home range scaling within a system might vary with competitive selection type. In instances where competition cannot be mitigated through relocation, high population densities would amplify selection for the most competitive behavioural and life history strategies (Milles et al. 2022). In contest competition dynamics, larger individuals are generally afforded a competitive advantage due to their physical superiority to smaller individuals (Fausch and White 1981). However, under scramble (i.e., exploitative) competition dynamics, where access to resources is mediated by number of competitors, individuals that grow more slowly (i.e., have lower BMR or SMR) have the advantage of decreased risk of starvation (Wilson 2014). Although the competitive dynamics of lemon sharks are not well described, lemon sharks display group living behaviour and show little evidence of conspecific aggression (Guttridge 2009), suggestive of scramble dynamics. In accordance, smaller, slower growing lemon sharks in the same population have highest survival rates during the first few years of life, indicating selection for conservative pace-of-life strategies (Dibattista et al. 2007, Dhellemmes et al. 2021a). As a result, conservative strategies likely predominate within this population, reducing the mean metabolic rate of the population, and as such, the population home range scaling rate.

## CONCLUSIONS

We provide the first direct test of the relationship between metabolism and home range allometry. Similarity of scaling rates in home range size and SMR confirmed a bioenergetic dependence of home range size. However, home ranges of lemon sharks scaled at lower rates than their SMR, highlighting considerable inconsistencies with the predictions of well-established models of resource competition leading to an increased home range scaling rate. Moreover, our results highlight the complexities of the relationship between competition and home range size, revealing that constraints on movement and the dynamics of competition experienced may determine whether the relationship is positive or negative. Selection for conservative life histories by scramble competition may only be apparent in situations where capacity for relocation is constrained by biological or environmental factors (e.g., predation risk, physiological or topographical barriers). This could be explored through comparative studies of the variation of individual survival under scramble competition dynamics across similar nursery habitats with differing levels of predation risk.

## ACKNOWLEDGEMENTS

The authors would like to thank Bimini Biological Field Station staff, S. Hart, C. Mason, A. Warrior, J. Whicheloe, and K. Yang, as well as interns who assisted with capturing and tagging of sharks. We thank the late Dr. S. Gruber for his dedication to furthering the understanding of elasmobranch ecology, and in generating a large body of foundational work on lemon shark physiology and ecology.

