## Supplemental files for "Intraspecific scaling of home range size and its bioenergetic dependence"

*The following supplement accompanies the article*

*Ecology*

---

**ELECTRONIC SUPPLEMENTARY INFORMATION**

**Table of contents** Page(s)

Fig S1: Histogram segregation plots 1
Fig S2: Temperatures experienced by individual sharks 2
Details of fitting RF models 3-4
Table S1: Variables used in RF models 5
Table S2: RF model performance 6
Fig S3: Variable importance for RF model (receivers 1-4) 7
Fig S4: Variable importance for RF model (receivers 5-8) 8
Fig S5: Variable importance for RF model (receivers 9-12) 9
Fig S6: Variable importance for RF model (receivers 13-16) 10
Supplementary References 11

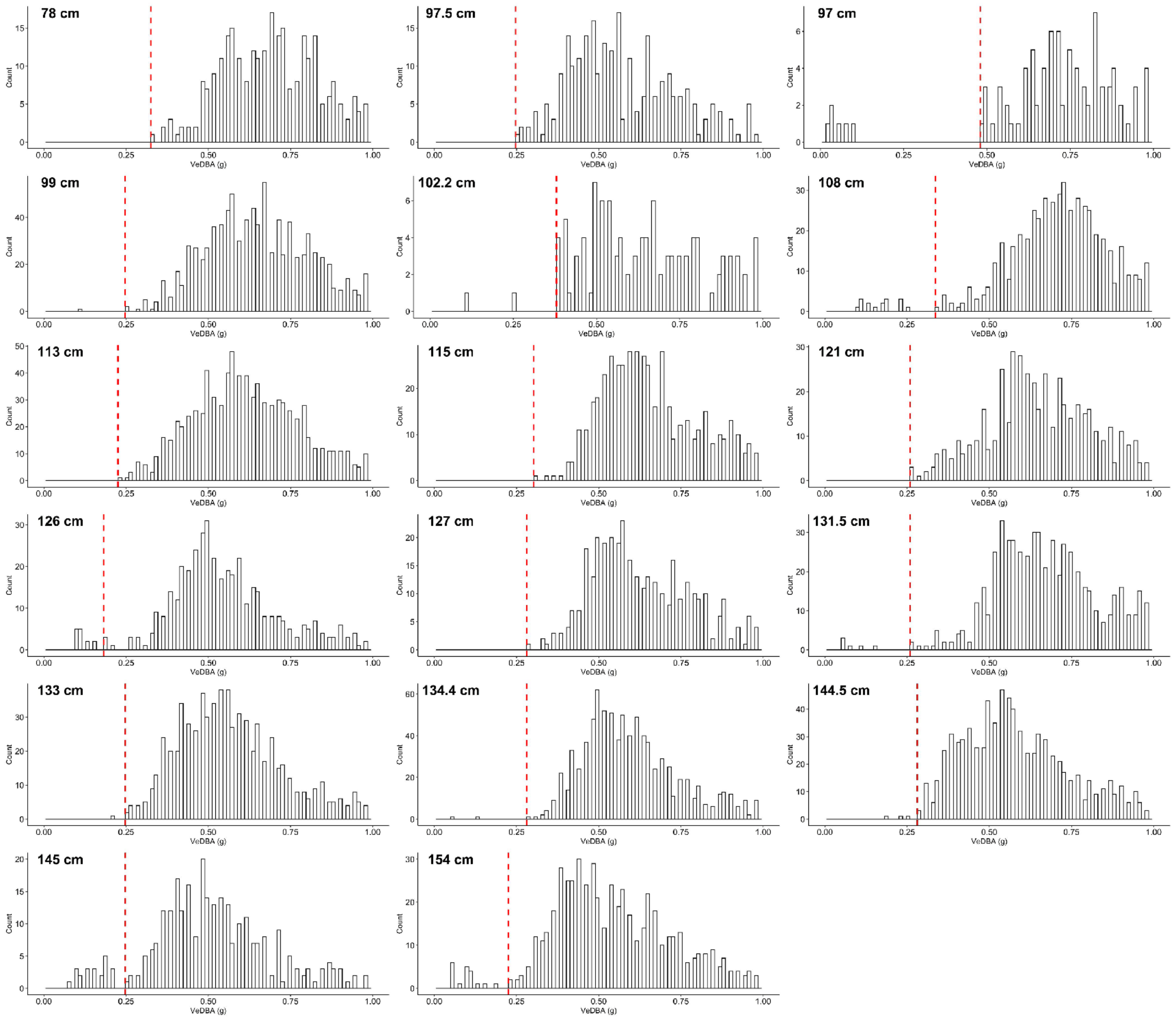


**Fig S1.** Histograms of VeDBA for each shark used in analysis. Red dotted vertical line indicates the value used to separate active from inactive VeDBA values for each shark, with values to the left of the line labelled as inactive and values to the right of the line labelled as active.

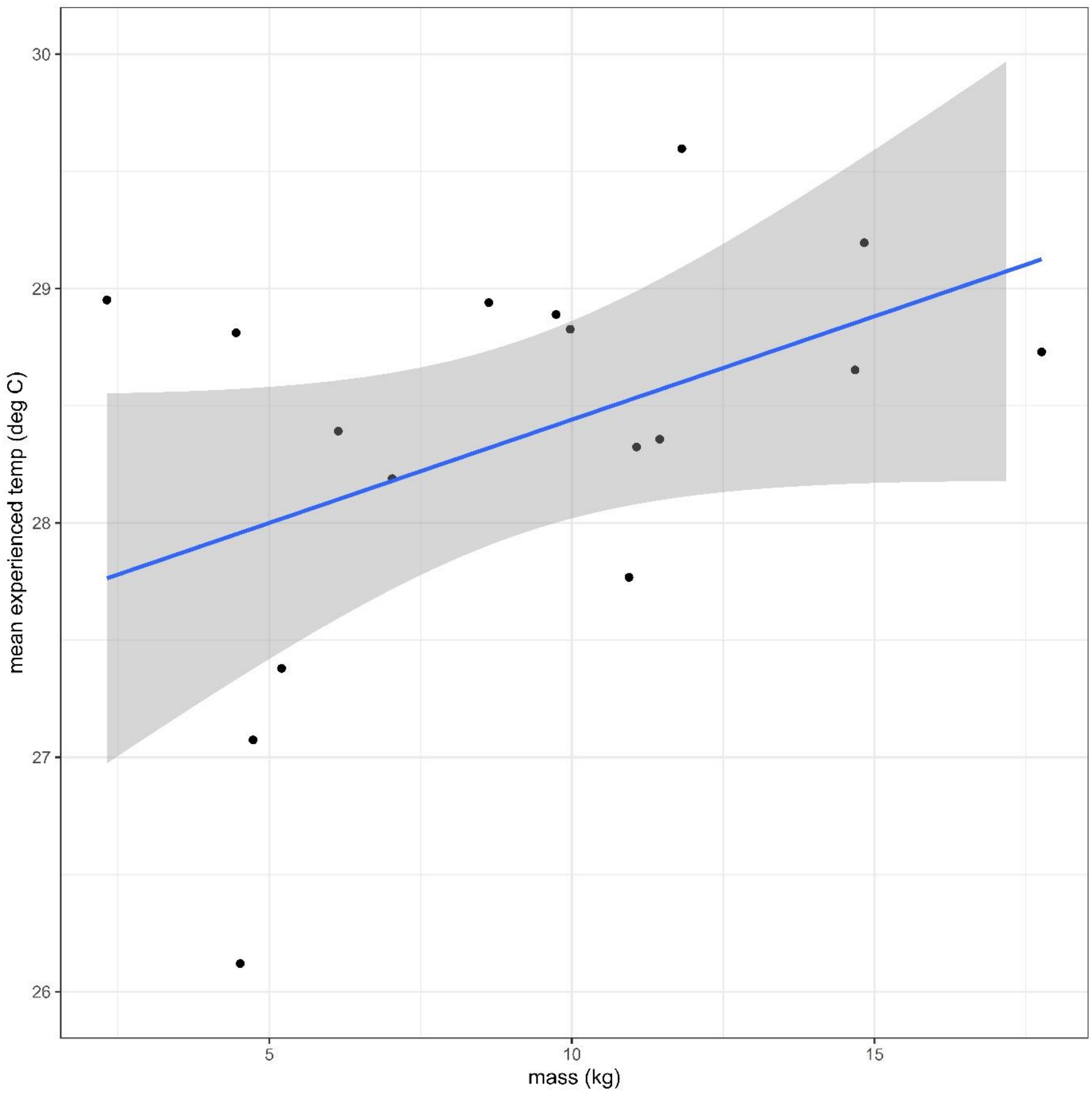


**Fig S2.** Mean daily temperature experienced by sharks plotted as a function of mass.

**Random Forest methods**

Multiple temperature loggers flooded or failed prematurely, so temperature data was either missing or was incomplete for many receiver locations. To reduce data incompleteness, we estimated water temperature using RF regression models, which has been shown to be an accurate method for environmental time-series forecasting [1]. A separate RF model was built for each receiver location where recorded water temperature data was available (n=16). For receivers with no available recorded water temperature data, temperatures were assumed to be the same as the closest receiver with similar habitat type and depth.

A random forest (RF) predict approach takes the aggregate prediction of multiple regression trees [2]. Each tree in the forest is created from a bootstrapped sample of training data for the total number of trees (ntree). The number of branches at each tree (mtry) is selected from a random subset of the input variables (p). New data is predicted by taking an average prediction of ntree regression trees. An internal estimate of error rate, mean square error (MSE), is calculated by predicting the data not contained at each bootstrap sample (out-of-bag, OOB; 36% of input data) and aggregating the OOB predictions.

All random forest regression analysis was performed in R (version 3.5.2). Models were trained using a dataset consisting of all observations of water temperature recorded by the respective temperature logger (°C) as the target (i.e., response) variable and a suite of corresponding environmental input (i.e., predictor) variables (Table S1). Where water temperature was unknown, the target variable did not contain data. RF models cannot predict from unordered categorical (factor) input variables. One-hot-encoding converts an unordered categorical vector to multiple binarised vectors where each binary vector of 1 and 0 indicates the presence of a class of the of the original vector. We used the one_hot function in the package mltools *[version 0.3.5; ,3]* one-hot encode tide phase.

Random forests were grown on the training dataset using the package randomForest *[version 4.6.14; ,4]*. We set mtry = 6 (default mtry = p/3) and ntree = 1000. For each receiver location, we predicted unknown water temperatures (°C) by fitting the trained model using the predict function in base R. We then assessed mean square error (MSE) as a measure of predictive performance for each RF regression model at each receiver location. RF models also allow for assessment of input variables importance for further interpretation, based on random variable selection for growing the RF (Fig. S3, S4, S5, S6). For each receiver location, variable importance was ranked by %IncMSE, indicating the increase of the MSE when the given input variable is randomly permuted. Overall, RF models for each receiver had a mean squared error ranging from 0.17 - 1.41 °C (mean: 0.67 °C; Table S2). All acoustic detections were matched with water temperature based on the temporally closest available water temperature for the respective receiver.

**Table S1.** Variables used for RF regression models to predict unknown water temperature for each receiver location.

| Variable type | Variable | Unit | Source† |
|---|---|---|---|
| target | Water temperature | °C | 1 |
| input | Day of the year | | 1 |
| | Hour of the day | | 1 |
| | Air temperature | °C | 2 |
| | wind direction | ° (0-360) | 2 |
| | Precipitation | mm | 2 |
| | Cloud cover | % | 2 |
| | Barometric pressure | bar | 2 |
| | Sun angle | ° (from horizon) | 3 |
| | Tide phase | High, Low, Rising High, Rising Low, Falling High, Falling Low | 4 |
| | Sea surface temperature | °C | 5 |
| | Lunar illumination | % | 6 |

†Data sources: 1) in-situ temperature loggers; 2) www.worldweatheronline.com; 3) *oce* package in R [5]; 4) www.tide.mobilegeographics.com; 5) www.seatemperature.info; 6) *lunar* package in R [6]

**Table S2.** RF regression model performance for each receiver location, indicated by OOB-MSE.

| REC # | OOB MSE |
|---|---|
| 1 | 0.80 |
| 2 | 1.61 |
| 3 | 0.28 |
| 4 | 0.27 |
| 5 | 1.41 |
| 6 | 0.43 |
| 7 | 0.17 |
| 8 | 0.26 |
| 9 | 0.18 |
| 10 | 1.07 |
| 11 | 0.33 |
| 12 | 0.72 |
| 13 | 0.46 |
| 14 | 1.20 |
| 15 | 1.09 |
| 16 | 0.37 |

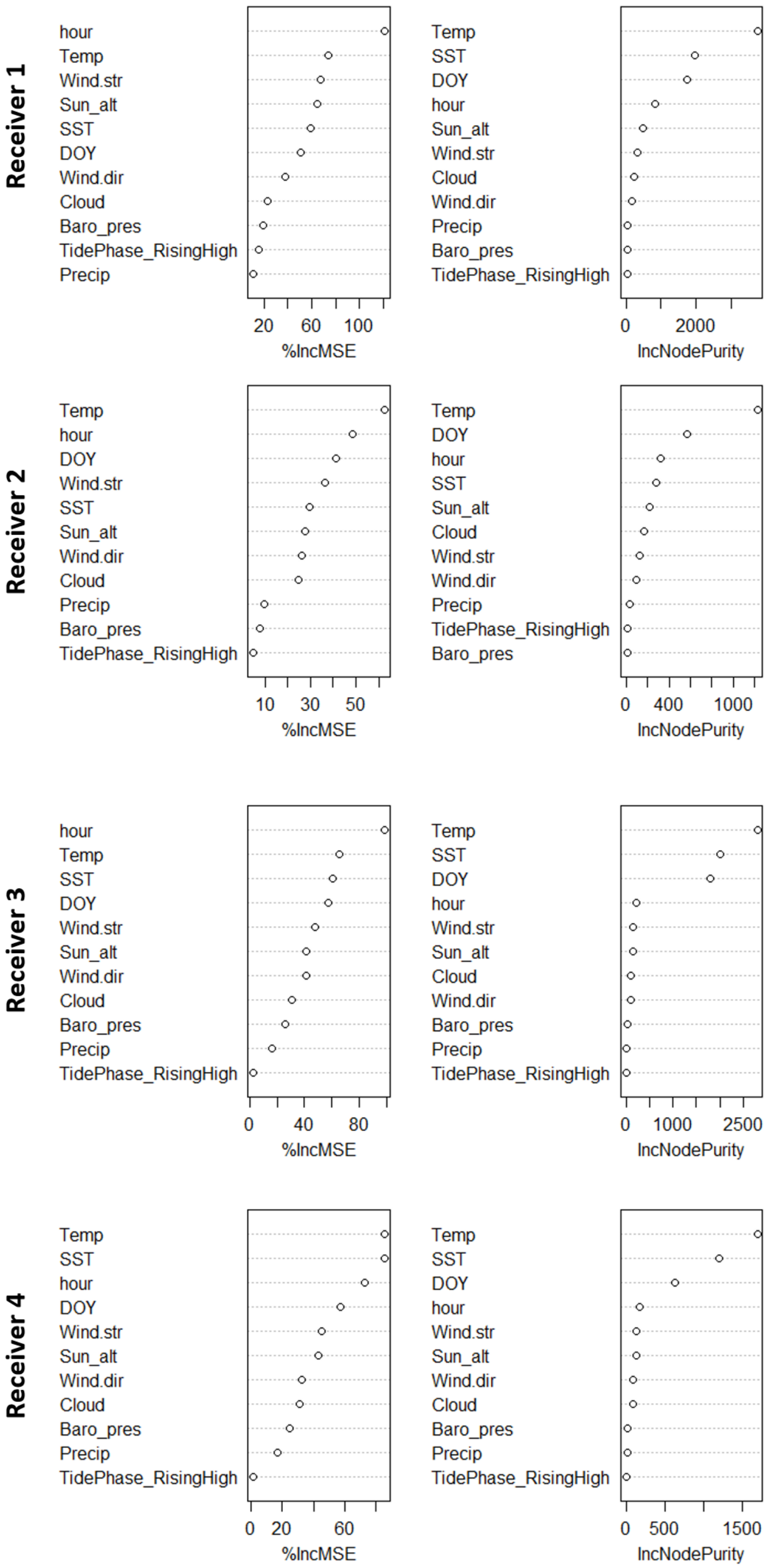


**Fig S3.** Variable importance within random forest models for receivers 1 to 4.

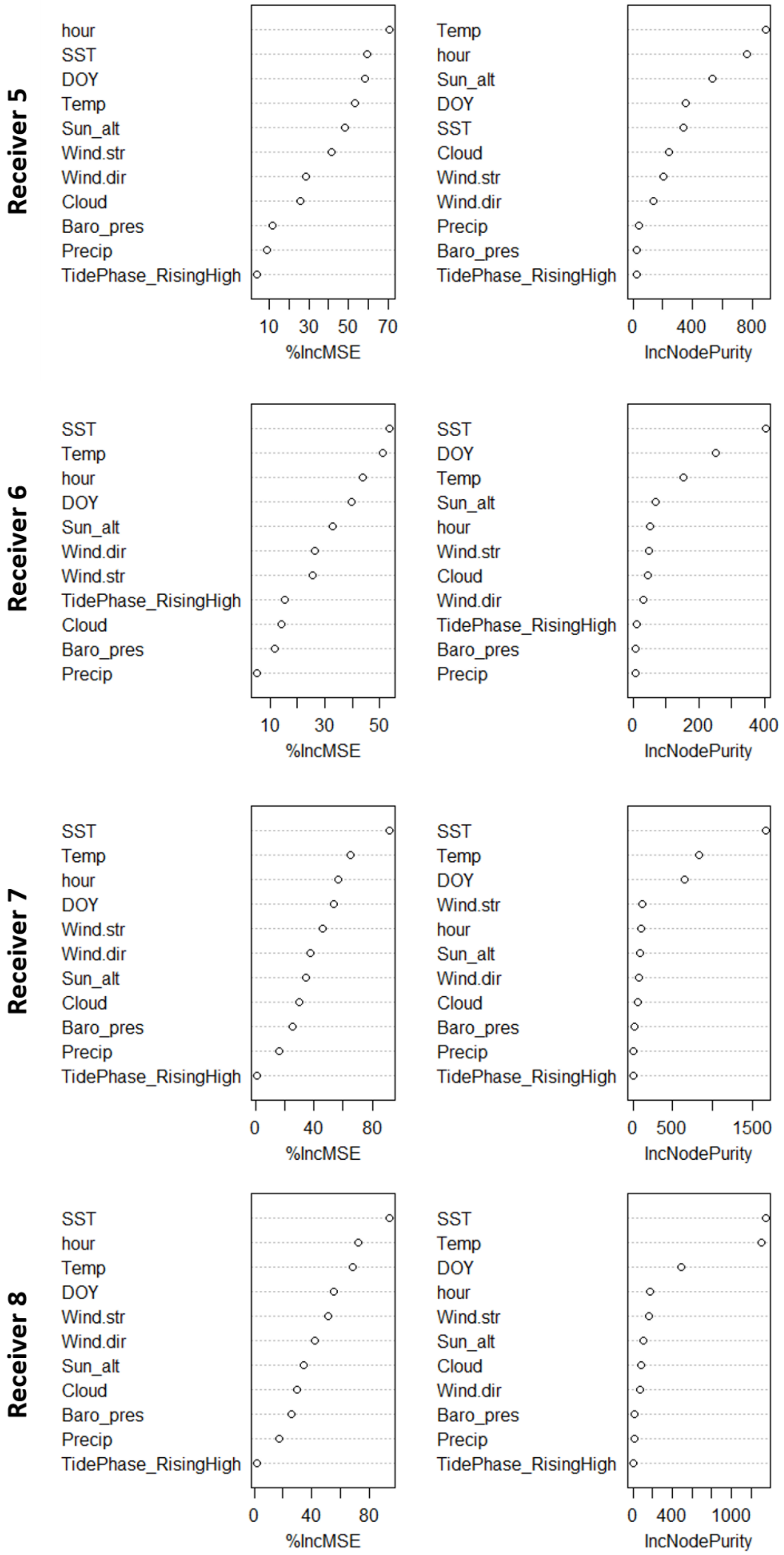


**Fig S4.** Variable importance within random forest models for receivers 5 to 8.

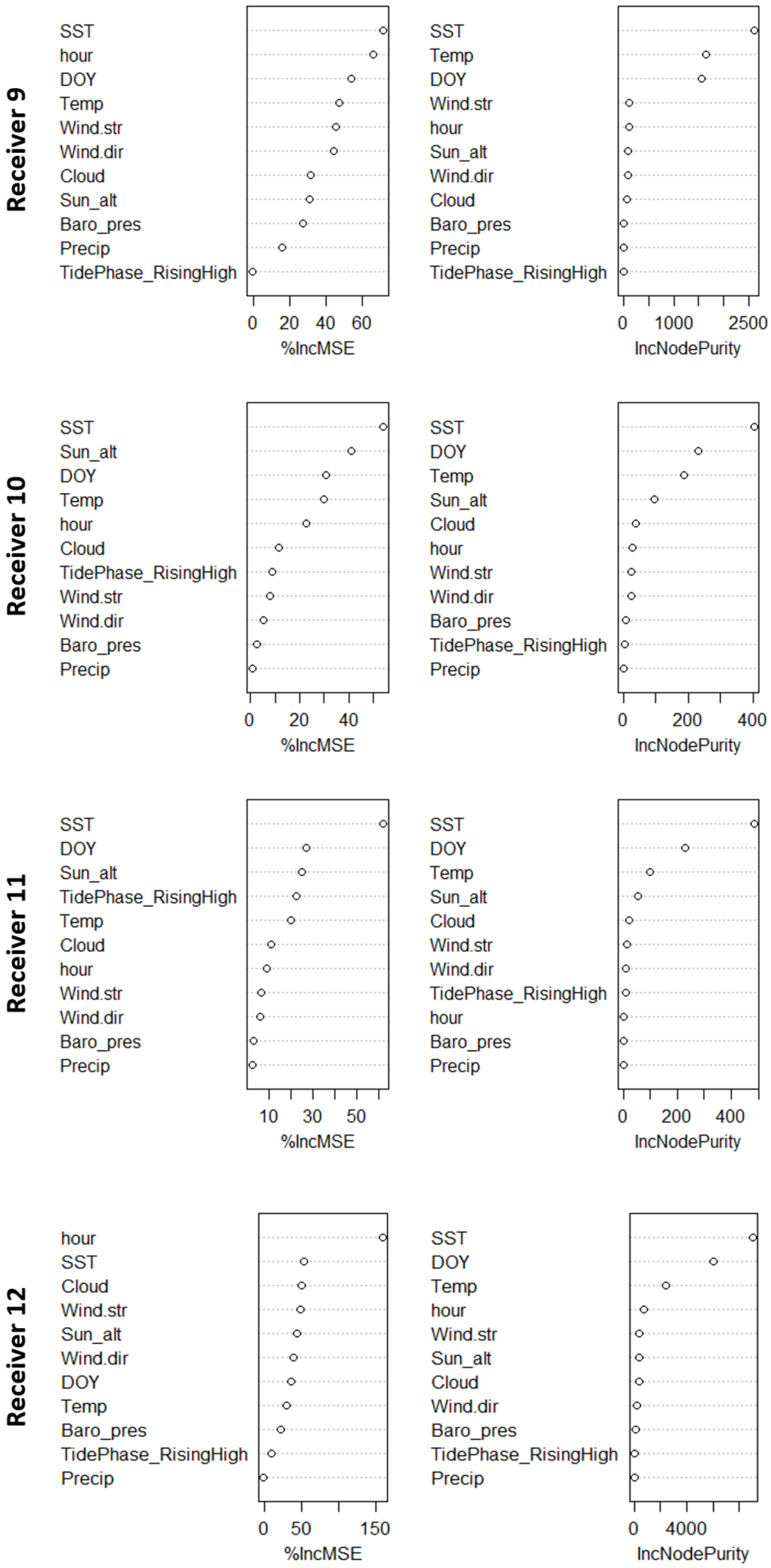


**Fig S5.** Variable importance within random forest models for receivers 9 to 12.

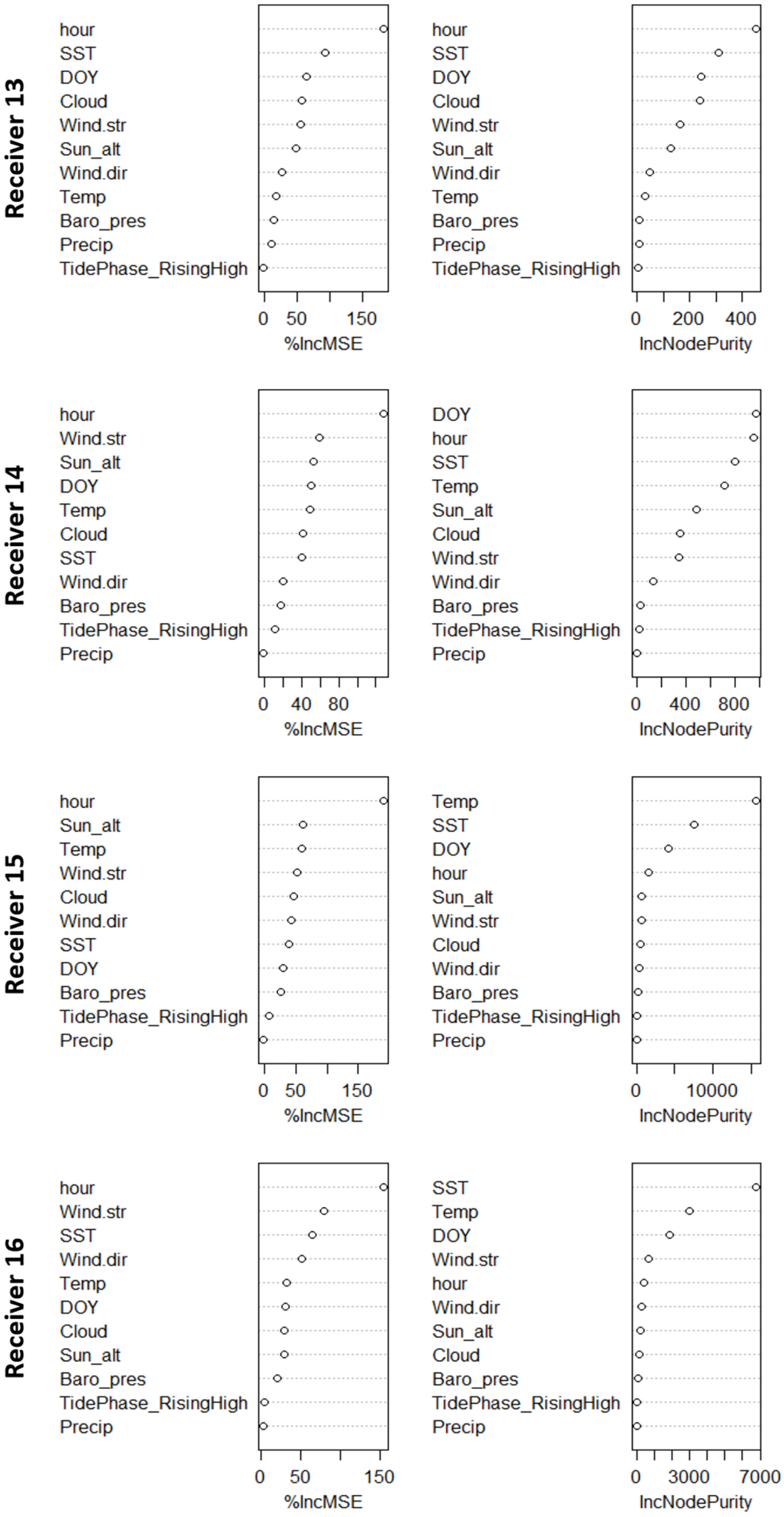


**Fig S6.** Variable importance within random forest models for receivers 13 to 16.
